# Investigation of cell nucleus heterogeneity

**DOI:** 10.1101/2020.07.08.193854

**Authors:** Noel Reynolds, Eoin McEvoy, Soham Ghosh, Juan Alberto Panadero Pérez, Corey P. Neu, Patrick McGarry

**Author notes:** Joint first authors (Noel Reynolds, Eoin McEvoy).

## Abstract

Nucleus deformation has been shown to play a key role in cell mechanotransduction and migration. Therefore, it is of wide interest to accurately characterize nucleus mechanical behavior. In this study we present the first computational investigation of the in-situ deformation of a heterogeneous cell nucleus. A novel methodology is developed to accurately reconstruct a three-dimensional finite element spatially heterogeneous model of a cell nucleus from confocal microscopy z-stack images of nuclei stained for nucleus DNA. The relationship between spatially heterogeneous distributions microscopic imaging-derived greyscale values, shear stiffness and resultant shear strain is explored through the incorporation of the reconstructed heterogeneous nucleus into a model of a chondrocyte embedded in a PCM and cartilage ECM. Externally applied shear deformation of the ECM is simulated and computed intra-nuclear strain distributions are directly compared to corresponding experimentally measured distributions. Simulations suggest that the nucleus is highly heterogeneous in terms of its mechanical behaviour, with a sigmoidal relationship between experimentally measure greyscale values and corresponding local shear moduli (*μ*_*n*_). Three distinct phases are identified within the nucleus: a low stiffness phase (0.17 *kPa* ≤ *μ*_*n*_ ≤ 0.63 *kPa*) corresponding to mRNA rich interchromatin regions; an intermediate stiffness phase (1.48 *kPa* ≤ *μ*_*n*_ ≤ 2.7 *kPa*) corresponding to euchromatin; a high stiffness phase (3.58 *kPa* ≤ *μ*_*n*_ ≤ 4.0 *kPa*) corresponding to heterochromatin. Our simulations indicate that disruption of the nucleus envelope associated with lamin-A/C depletion significantly increases nucleus strain in regions of low DNA concentration. A phenotypic shift of chondrocytes to fibroblast-like cells, a signature for osteoarthritic cartilage, results in a 35% increase in peak nucleus strain compared to control. The findings of this study may have broad implications for the current understanding of the role of nucleus deformation in cell mechanotransduction.

## Introduction

The morphology and deformation of cellular nuclei influences differentiation, immune response, migration and disease development (1). Regarding migration, many cancer and immune cells have highly deformable nucleus (2-5), increasing their migratory potential by enabling passage through narrow matrix pores. In relation to cell differentiation, a recent study has shown that an increase in matrix stiffness can induce an increase in MSC contractility, which tenses the nucleus to favor lamin accumulation in the nuclear envelope and results in osteogenesis over adipogenesis (6). Others have identified that MSC nuclear morphology depends on cell density, becoming highly rounded at high densities, leading to an increased expression of genes typical of pre-, peri-, and post-chromatin condensation events (7). Charlier et al. reported that a population of cells in cartilage during osteoarthritis (OA) display a similar expression profile to dedifferentiated chondrocytes in vitro (8), suggesting that a subset of mature chondrocytes on OA undergo trans-differentiation to a fibroblast-like phenotype with a different deformation state. Therefore, the development of computational models that accurately predict nucleus deformation would represent a significant advance in current understanding of the link between nucleus mechanical deformation and the biological function of cells, potentially informing strategies for control of migration, differentiation, and engineering of functional tissue.

Biomechanical studies to date have considered the nucleus to be homogeneous and generally stiffer than the surrounding cytoplasm (9-17). Parallel plate compression studies, in which material properties of the components of isolated cells are determined through inverse finite element analysis of experiments (16, 18), suggest that nuclei near incompressible with shear moduli in the range from ∼1kPa to ∼3 kPa. However, analysis of micropipette aspiration studies consistently report lower values of shear modulus lower than this range (Deguchi et al. (15), Guilak et al. (9), and Zhao et al. (19)). Such micropipette experiments are typically performed on suspended cells, in contrast to adhered spread cells used for confined compression experiments. However, using an active model for cytoskeletal evolution and contractility during spreading, combined with a new experimental methodology for micropipette aspiration of spread cells, (20) report a nucleus shear modulus of only 0.07 kPa, again with near incompressibility being observed. Micropipette aspiration of isolated chondrocyte nuclei by (21) report a shear modulus of 0.008 kPa. The vast differences between reported values of the apparent nucleus shear modulus for compression and micropipette experiments suggests that the mechanical deformation of the nucleus is dependent on the applied mode of deformation, and it is not accurately predicted by a simplistic assumption of homogeneous material behaviour. T Several studies suggest that the cell nucleus is elastic with fully recoverable deformation following the application of moderate to large deformations (18, 22, 23). However, Pajerowski et al. (24) report that permanent viscoplastic deformation of the nucleus can occur following knockout of lamin A/C, and Thiam et al. (25) report rupture of the lamin following cell migration through narrow channels. In an experimental investigation by Henderson et al. (26), 3D strain distributions inside the nuclei of single living cells embedded within their native extracellular matrix (ECM) were determined during external application of tissue shear deformation. During deformation of a cartilage tissue explant, strain is transferred to individual cell nuclei, resulting in submicron displacements. Local deformation gradients were determined from confocal images of nuclear DNA distributions before and after the application of an applied shear loading were used to determine three-dimensional intra-nuclear distribution of strain. Shear strain localisations in some regions of the nucleus were shown to be five-fold higher than the that in the ECM.

In the current study, we investigate the role of intra-nuclear material heterogeneity in the intra-nuclear strain magnification. We develop a novel modelling approach to construct a heterogeneous finite element model of the chondrocyte nucleus based on greyscale values obtained from confocal z-stacks of the DNA in nuclei within cartilage tissues explants (26). We construct an RVE model of a cartilage explant in which, in addition to the heterogeneous nucleus, we include the chondrocyte cytoplasm and actin cytoskeleton (27), the pericellular matrix (PCM) and the ECM (28). By comparing computed intra-nuclear heterogeneous strain distributions to experimental measurements, we explore the relationship between heterogeneous nucleus shear moduli and corresponding local greyscale values (which are dependent on local DNA concentration). We also explore the influence of the nucleus envelope on heterogeneous nucleus strain. Additionally, we explore the influence of a phenotypic shift of chondrocytes to fibroblast-like cells (a signature of osteoarthritis (8)) on intra-nucleus strain distribution.

## Model development

### A constitutive model for the mechanical behavior of chondrocytes embedded in cartilage

We develop a representative volume element (RVE) for cartilage tissue (Fig. 1), in accordance with the methodology proposed by Dowling et al. (28). Briefly, an RVE comprising of a chondrocyte cell surrounded by a peri-cellular matrix (PCM) embedded in a cuboidal ECM is modelled. The cuboid has a side dimension of 60 μm, based on observed cell spacing in situ. The nucleus, cell, and PCM are assumed to be spherical with diameters of 7.5 μm, 16 μm, and 22 μm, respectively, again based on experimental observation. It should be noted that dimensions are chosen such that the volume fraction of cells and ECM is representative of cartilage tissue (26). To replicate the application of 15% shear strain to the cartilage explants in the experiments of, displacement boundary conditions are applied to the upper surface of the RVE model. The bottom surface of the cartilage explant and the RVE are rigidly fixed in all directions. Images of the tissue explant, RVE, and nuclei before and after shear deformation are shown in Fig. 1.

**Figure 1:**
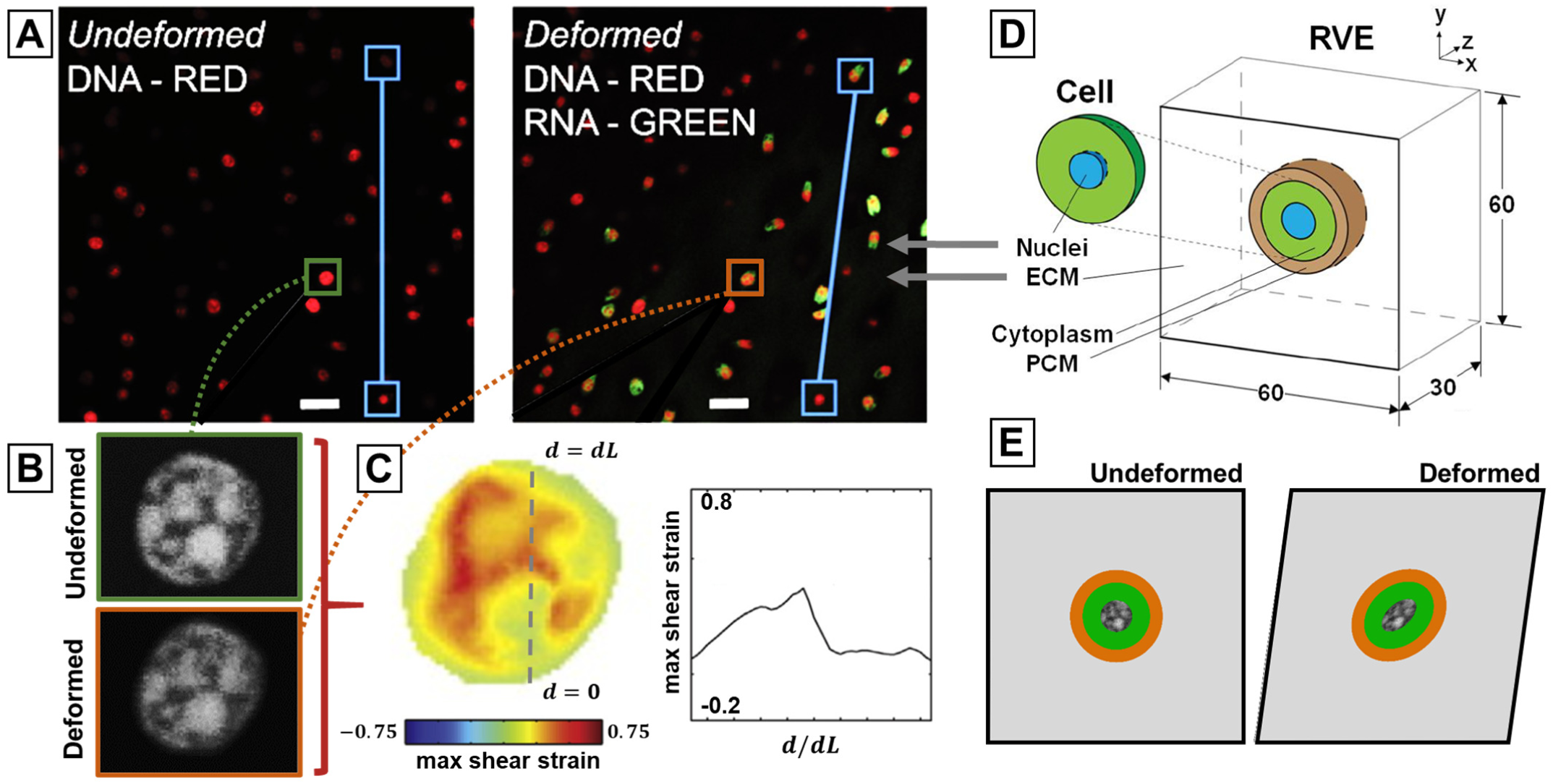
(A) Undeformed and deformed confocal image of cartilage tissue explant showing DNA in red and nascent RNA in green as experimentally observed by Henderson et al. (26). The blue line indicates the relative motion between two nuclei in the explant before and after shear deformation. Scale bar = 20 μm. (B) Greyscale confocal image of a DNA in a single nucleus before and after shear deformation. (C) Shear stress along nucleus section. (D) Schematic of an RVE of cartilage consisting of a single chondrocyte cell surrounded by a PCM embedded in a cuboidal ECM. (E) Mid-section of the RVE before and after shear deformation.

For full details on material properties of the ECM, PCM, and cell cytoplasm the reader is referred to the study of Dowling et al. (28). Briefly, the ECM and PCM are modelled using an isotropic Neo-Hookean hyperelastic constitutive formulation with a Cauchy stress tensor given as:

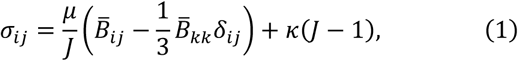

where *μ* and *κ* are material shear and bulk moduli, respectively, *J* is the volumetric Jacobian, and 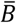 is the isochoric left Cauchy Green deformation tensor. Following from the work of Dowling et al. (28). The shear and bulk moduli of the ECM (PCM) are 400 kPa (22 kPa) and 2.0 MPa (100 kPa), respectively. Preliminary analyses reveal that addition of an anisotropic hyperelastic component (29), representing collagen fibre distributions in the cartilage ECM, does not significantly influence the response of the RVE to the applied shear deformation of Henderson et al. (26). In-vitro mechanical testing of isolated chondrocytes by Dowling et al. (27) reveals that the actin cytoskeleton is the dominant component in the shear response of chondrocytes. Furthermore, computational simulation of in-vitro experiments revealed that the actin cytoskeleton’s mechanical contribution to chondrocyte shear resistance is not described by a standard hyperelastic formulation, but rather is described by an active bio-chemo-mechanical constitutive law that incorporates active Hill-type contractility and tension dependent remodelling (27). Briefly, based on the formulation of Deshpande et al. (30) and the implementation by McGarry et al. (31) and Ronan et al. (32), a first order kinetic equation is used to capture formation and dissociation of the actin cytoskeleton:

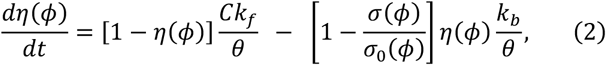

where *η*(*ϕ*) is the non-dimensional activation level of a stress fibre in the *ϕ* direction within the actin cytoekeleton. *k*_*f*_ and *k*_*b*_ are forward and backward reaction rate constants, respectively. *C* is an activation signal for SF formation that decays over time (*C* = exp(−*t*/*θ*)) where *θ* is a decay constant for the signal. To simulate the contractile behaviour of the fibre bundle a Hill-like equation is used:

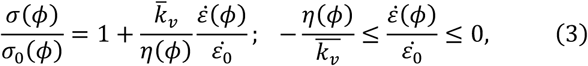

where 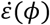 is the strain rate of a stress fibre in direction *ϕ* within the actin cytoskeleton. Actively generated tension decreases if 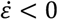. Under steady state conditions 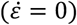, or during extension 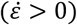 an isometric tension level (σ_0_ = *η*σ_*max*_) is generated. 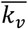 and 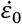 re model parameters. Dowling et al. (28) also showed that the mechanically passive components of the cell, including the cytoplasm, microtubules and intermediate filaments, are accurately represented by placing a passive neo-Hookean hyperelastic component in parallel with the active actin cytoskeleton component. The use of this active remodelling and contractility formulation for chondrocytes is a key feature of our model. An early study by McGarry and McHugh (33) demonstrated that the apparent mechanical properties of chondrocytes change with increasing levels of cell spreading. Dowling and McGarry (34) showed the active framework captures the key relationship between cell deformability and morphology, correctly predicting stiffer shear behaviour for more spread morphologies. The active framework is implemented in a *user-defined material subroutine* (*umat*) in the finite element software Abaqus.

Parameters for the active modelling framework are confined to previously reported values (27) determined from chondrocyte shear loading. Briefly, σ_*max*_ = 0.85 *kPa*, 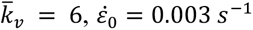, *θ* = 70*s*, *k*_*b*_ = 10, *k*_*f*_ = 1, *μ*_*cyto*_ = 0.54 *kPa* and *κ*_*cyto*_ = 2.5 *kPa*. In the current study we use a novel approach to construct and analyse a mechanically heterogeneous, as described in the following section). Each material within the nucleus is assumed to be hyperelastic (via Eqn. 1), such that all nuclear deformations are fully recoverable.

### Development of a novel heterogeneous nucleus finite element model

The eight z-stacks of a confocal image of a single nucleus stained for nucleus DNA taken from the study of Henderson et al. (26) are shown in Fig. 2A. The image is taken from a cell in-situ in the undeformed cartilage explant. Each of the experimentally obtained z-stacks shown in Fig. 2A have a resolution of 50×50 pixels.

**Figure 2:**
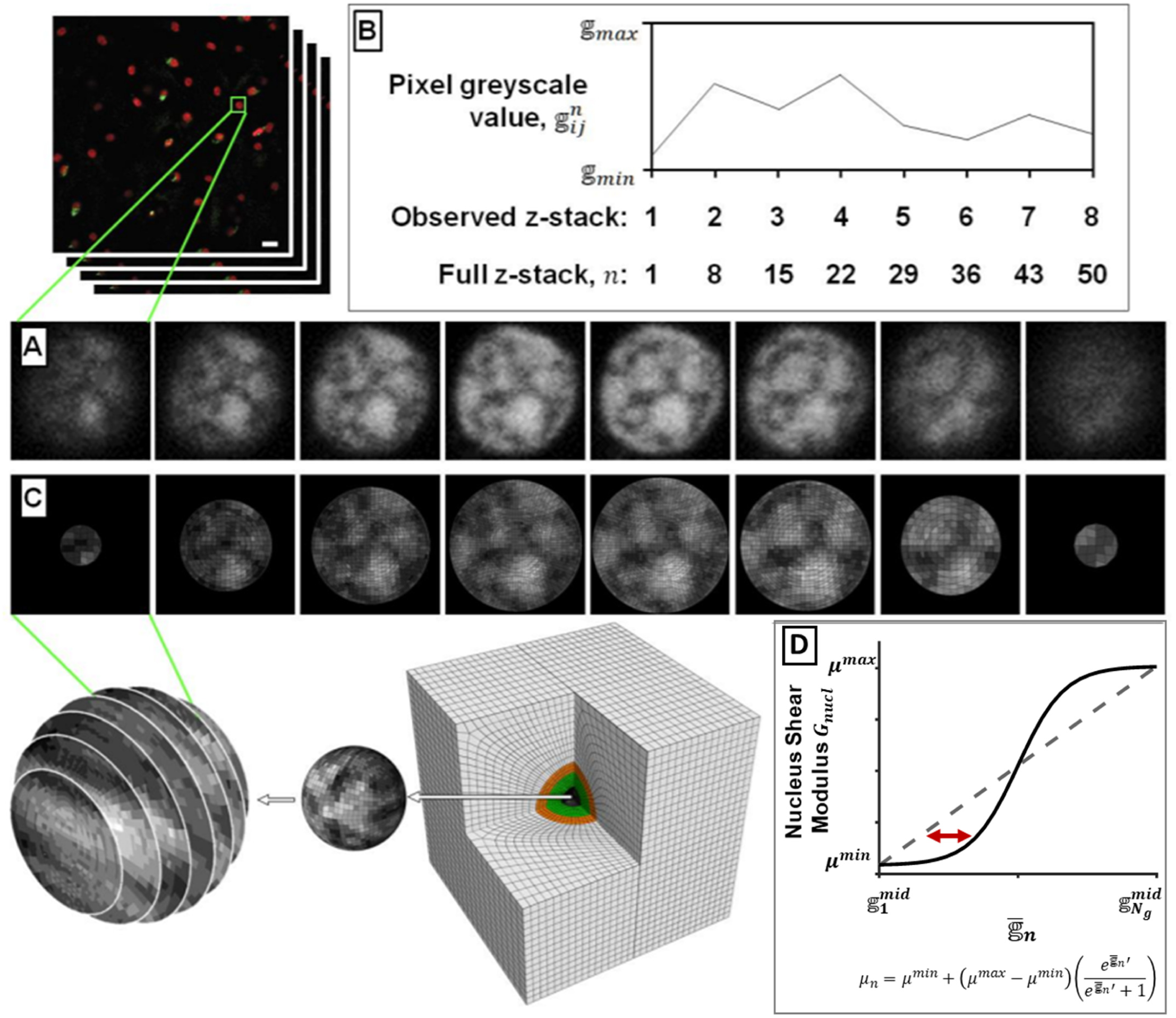
(A) A representation of image stacks of a cartilage tissue explant. Greyscale images of a single nucleus from each confocal z-stack showing DNA intensity as experimentally observed by Henderson et al. (26). Scale bar = 20 μm. (B) Graphical representation of linear interpolation between observed greyscale values. (C) Corresponding slices of the nucleus in the finite element RVE. The contour plot illustrates assigned material sections ranging from stiff (light) to compliant (dark). (D) Graphical representation of shear modulus assignment to each group.

In the current study, a finite element model of the undeformed nucleus is constructed as follows: The spacing between adjacent z-stacks (∼1 *μm*) is equivalent to the length of seven pixels. In order to construct a cube of 50×50×50 uniformly distributed greyscale values, six additional “model” z-stacks are generated between each pair of adjacent experimental z-stacks by using linear interpolation between the grayscale values of corresponding pixels. For example, taking the experimental grayscale values 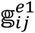 and 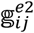 at pixel at position *i, j* in the 50×50 image for experimental z-stack 1 and z-stack 2, respectively, we use linear interpolation to determine 6 intermediate “model” grayscale values 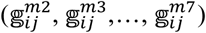. Repeating this process for all positions *i, j* results in the construction of six “model” z-stacks between the adjacent experimental z-stacks. This results in a total of 50 uniformly distributed grayscale values in the z-direction of the cube that contains the nucleus, along with 50 uniformly distributed grayscale values in the x- and y-directions of the cube.

A finite element mesh for the spherical nucleus is created using the commercial finite element software Abaqus (DS Simulia, RI, USA). By assuming the diameter of the spherical nucleus is equal to the edge dimension of the 50×50×50 cube of pixel grayscale intensities, a mesh density is chosen such that the number of finite elements per unit volume is equal to the number of pixels per unit volume. The 3D coordinate of the centroid of each element is associated with a corresponding greyscale value in the 3D cube of pixel intensities. Elements are assembled into *N*_*G*_ greyscale groups (GSGs) based on greyscale values; (in the current study *N*_*G*_ = 10). For example, grayscale group 1 (GSG1) contains all elements with greyscale values between 𝕘^*min*^ and 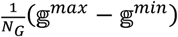, where 𝕘^*max*^ and 𝕘^*min*^ are the maximum and minimum greyscale values found in the entire domain, respectively. *E*_*ij*_, an element at location (i,j), is assigned to *GSGn* based on the following criterion:

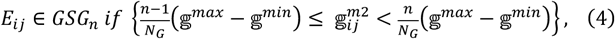

where *n* = 1, *N*_*G*_. The mid-range grayscale value of *GSGn* is given as 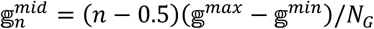. It is also convenient to define a non-dimensional normalised mid-range grayscale value, 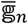, for each *GSG*, such that

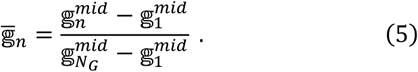

The pseudo z-stack generation, centroid coordinate calculations, greyscale mapping, and element grouping assignments are performed using scripts written MATLAB (The Mathworks, Natick, MA). To compare the greyscale map in the finite element mesh to experimentally observed z-stacks, each group in the finite element mesh of the nucleus is assigned a greyscale value so that colour ranges from black for GSG1 to white for GSG10. The planes shown in Fig. 2C are chosen to correspond with the experimentally observed z-stacks in Fig. 2A. This analysis demonstrates that the complex patterns of greyscale distributions observed experimentally are accurately replicated through our model construction protocols. As illustrated in Fig. 2D, the shear modulus *μ*_*n*_ of GSGn is related to the grayscale value 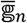 through the following sigmoidal relationship:

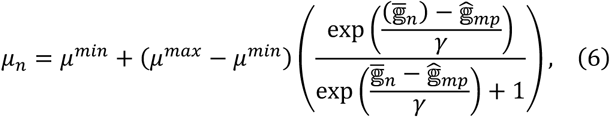

where the parameters *μ*^*max*^ and *μ*^*min*^ correspond to the minimum and maximum location specific shear moduli in the nucleus, respectively, such that *μ*_1_ = *μ*^*min*^ and *μ* _*NG*_ = *μ*^*max*^. The parameters 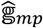 and γ set the mid-point and width of the sigmoidal distribution, respectively. The function collapses to a linear distribution of shear modulus as a function of 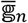 for the case of *γ* → ∞. This linear distribution is also shown in Fig. 2D. In addition to exploring sigmoidal and linear distributions of *μ*_*n*_, we also demonstrate the significant inaccuracies in the computed nucleus strain state if homogeneous properties are assumed. To enforce condition of near incompressibility throughout the nucleus, based on the findings of Reynolds et al. (20) and Weafer et al. (23), we assume a ratio of bulk modulus to shear modulus (*κ*_*n*_/*μ*_*n*_) of ∼30 for all GSGs.

## Results

### Nuclear stiffness is heterogeneous with distinct phases associated with high and low DNA content

A finite element parametric investigation was performed to identify the relationship between greyscale value (DNA concentration) and shear modulus in a heterogeneous nucleus that results in the best-fit prediction of shear strain along a linear path through the centre of the nucleus, as measured experimentally by Henderson et al. (26). Specifically, a sigmoidal relationship between 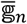 and *μ*_*n*_ was explored across the following parameters ranges: 0.08 *kPa* ≤ *μ* ^*max*^ ≤ 8 *kPa*; 0.08 *kPa* ≤ *μ* ^*min*^ ≤ 8 *kPa*; 0.2 ≤ 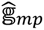 ≤ 0.8; 0.015 ≤ *γ* ≤ 0.15. Fig. 3 presents the best-fit 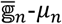 relationship (shown in Fig. 2D) computed for the parameters: *μ*^*max*^ = 4 *kPa, μ*^*min*^ = 4 *kPa*, 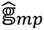 = 0.5; *γ* = 0.083, with a comparison between the computed and experimental distribution of shear strain through the centre of the nucleus shown in Fig. 3A. Computed results exhibit a peak strain (0.375) similar to the experimental value (0.374). The location of this peak (*d/d*_*L*_=0.536) is reasonably close to the experimental location (*d/d*_*L*_=0.471). Additionally, computed magnitudes (∼0.09) and locations (0.55 < *d/d*_*L*_ *< 1*.*0*) of low strain regions are reasonably similar to corresponding experimental values. Contour plots of the distribution of maximum shear strain on a plane through the centre of the nucleus are shown in Fig. 3B. Experimental and computed distributions exhibit similar level of contrast between high and low strain regions. Importantly, the inclusion of a nuclear envelope in the computational model results in a low strain region on the periphery of the nucleus similar to that measured experimentally. The computed distribution of shear strain in each GSG is presented in Fig. 3C. Interestingly, for low greyscale (low DNA concentration) regions, the position of a material point in the nucleus is an important indicator of the strain level. For example, material points in GSG1, GSG2, GSG3 all have similarly low shear moduli (∼0.04*μ*^*max*^) due to the sigmoidal-type distribution. However, low strains occur throughout GSG1 and GSG2 because all elements are close to the stiff nucleus envelope, which provides a deformation shielding effect. Elements in the interior of GSG3 exhibit high strains, but elements near the nuclear envelope again exhibit low strains. A similar pattern is also observed in regions of medium greyscale (DNA concentration) values, i.e. GSG4 (*μ*_4_ = 0.1*μ*^*max*^) and GSG5 (*μ*_5_ = 0.2*μ*^*max*^). The relative high greyscale (DNA concentration) regions (GSG6-GSG10) exhibit very low levels of deformation, despite the fact that the majority of such material points are located in the interior of the nucleus. Fig. 3F shows the computed distribution of shear modulus as a function of material volume. The low stiffness (low DNA concentration) GSG1-4 regions (0.17 ≤ *μ* _*n*_ ≤ 0.63) comprise ∼57% of the nucleus volume. In contrast, the high stiffness (high DNA concentration) regions GSG7-10 (1.48 ≤ *μ*_*n*_ ≤ 2.7) comprise only ∼12% of the nucleus volume. We label GSG5-6 as the intermediate stiffness region (3.58 ≤ *μ*_*n*_ ≤ 47), comprising of ∼31% of the total volume. This suggests that low DNA (high RNA) regions make up the majority of the nucleus volume, but that the nucleus cannot be considered as a bi-modal structure, with 30% of the volume exhibiting an intermediate stiffness with moderate DNA concentrations.

**Figure 3:**
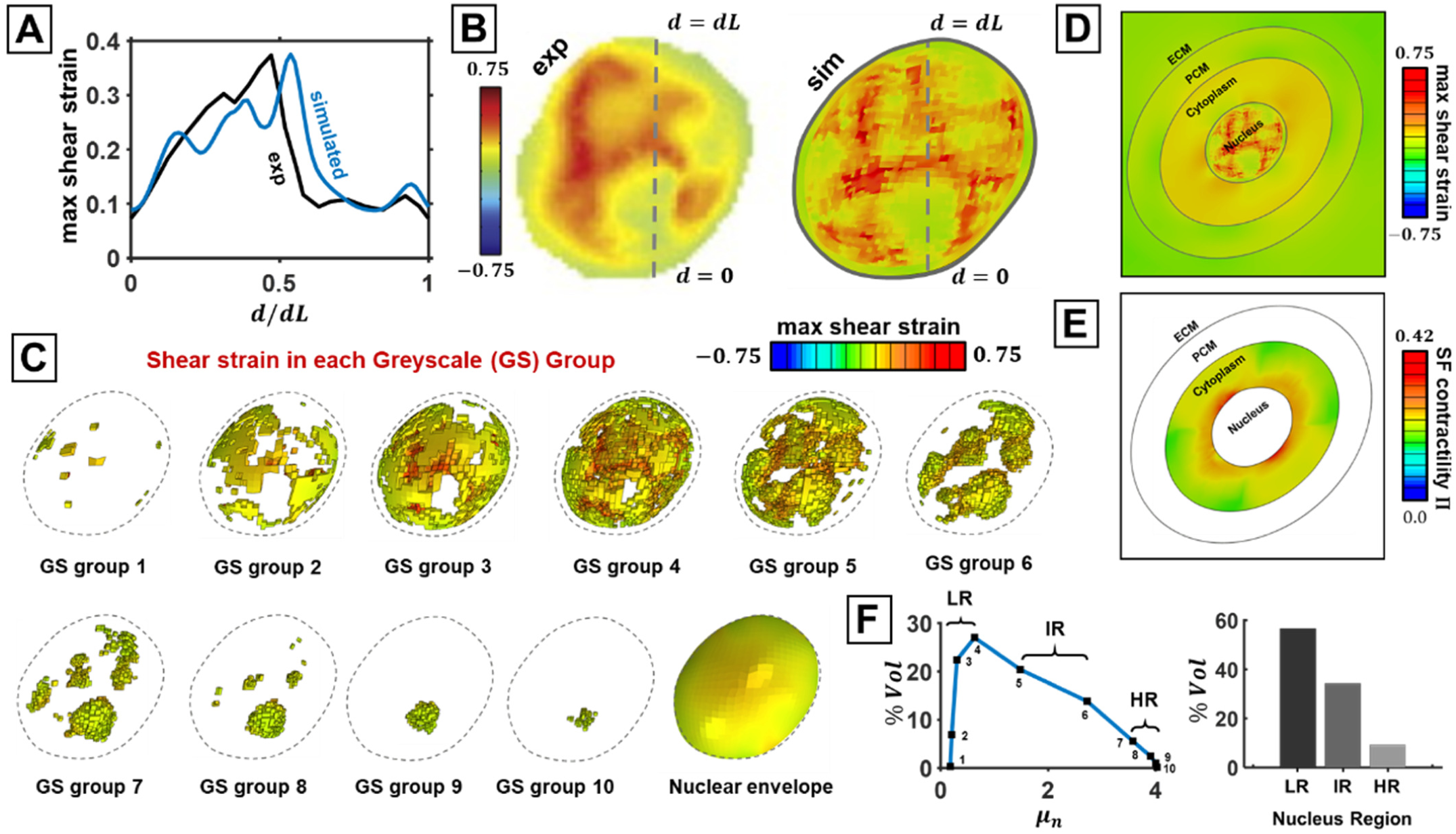
(A) Experimental and predicted shear strain along nucleus section following deformation of a heterogeneously stiff nucleus; (B) Experimental and simulated contour plots of shear strain in nuclear mid-section; (C) 3D separation of Greyscale groups (GSG) with associated predicted shear strain; (D) Shear strain across tissue model showing distinct regions for the ECM, PCM, cytoplasm, and nucleus; (E) Active actomyosin contractility Π = *η*_*max*_ − *η*_*avg*_ in response to applied loading; (F) Nuclear volume of each GSG as a function of associated shear modulus. Three distinct regions are identified: High DNA (HR), Intermediate (IR) and Low DNA (LR).

### The nuclear envelope acts as a strain shield

The distribution of strain is predicted to be non-uniform across the cell cytoplasm, PCM and ECM (Fig. 3D). The strain is lower in the cytoplasm than in the PCM or ECM, as expected. However, the cytoplasm strain is significantly higher than the strain in high DNA regions of the nucleus and is significantly lower than the strains in the low DNA regions of the nucleus. This demonstrates the common assumption that the nucleus acts a stiff low deformation component is not accurate and suggests that alterations in cytoplasm deformation and contractility may significantly alter localised deformation within the nucleus. A previous experimental-computational study by Dowling et al. (28) demonstrated that the actin cytoskeleton contractility is the key contributor to the shear resistance of chondrocytes. In Fig. 4A we remove the nuclear envelope from the model to simulate the effect of lamin A/C depletion on nucleus shear strain distribution. Strains are significantly increased (∼4-fold) in low DNA regions near the periphery of the nucleus. High DNA regions in the interior of the nucleus are not strongly influenced by removal of the nuclear envelope.

**Figure 4:**
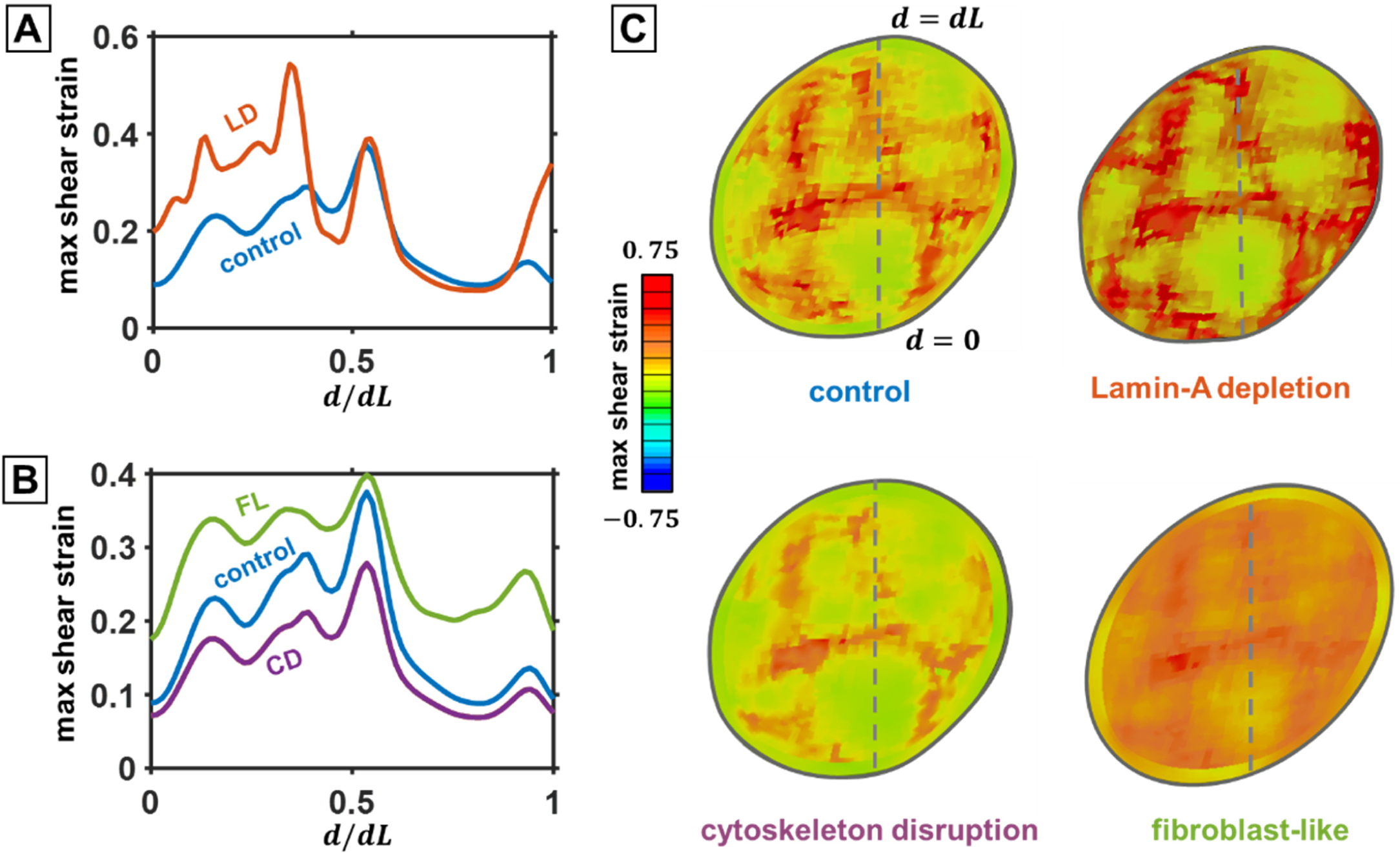
(A) Predicted shear strain along nucleus section following deformation of a heterogeneously stiff nucleus under control conditions and with a disrupted nuclear envelope; (B) Predicted shear strain along nucleus section following deformation of a heterogeneously stiff nucleus under control conditions and with a disrupted or fibroblast-like actomyosin network; (C) Associated contour plot across nuclear mid-section.

### Dedifferentiation of chondrocytes to fibroblast-like cells increases nuclear strain

In Fig. 4B-C we explore the influence of cytoskeletal suppression on nucleus deformation by reducing the contractility parameter σ_*max*_ to zero. This is predicted to reduce nucleus shear strain by ∼0.06 in low DNA regions and by ∼.02 in high DNA regions. In contrast, a phenotypic shift of chondrocytes to fibroblast-like cells, a signature for osteoarthritic cartilage, is simulated by increasing the contractility of the actin cytoskeleton to a level associated with fibroblasts. Peak nucleus strains increase by 35% compared to control as the nucleus becomes more ellipsoidal. Our simulations therefore suggest that de-differentiation to a fibroblast-like phenotype significantly elevates heterogeneous intra-nuclear strain, potentially altering cell function. This result is also supported by recent work from Alisafaei et al. (35) that revealed how alterations in nuclear shape and associated gene expression can be driven by cytoskeletal contractility. The predictions of the current study further advance these findings by demonstrating that impairment of actomyosin force generation reduces the shear strain in low DNA regions of the nucleus by ∼26%.

### A linear relationship between greyscale and shear modulus is inaccurate and a homogeneous nucleus is highly inaccurate

Finally, we demonstrate that the assumption of a linear relationship between greyscale 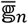 and shear modulus *μ*_*n*_ does not provide an accurate prediction of the large differences in strain between low and high DNA regions (Fig. 5A-B). While the parameter *μ*^*max*^can be chosen to accurately predict the low strain in high DNA regions, successive reductions of the value of *μ*^*max*^results in a plateau in the computed maximum strain at a value that is ∼50% lower than the observed experimental value. Further, we highlight the significant inaccuracy that results from the assumption of a homogeneous nucleus (Fig. 5C-D). Neither a high nor low value of shear modulus provides a reasonable prediction of the true distribution of deformation throughout the nucleus.

**Figure 5:**
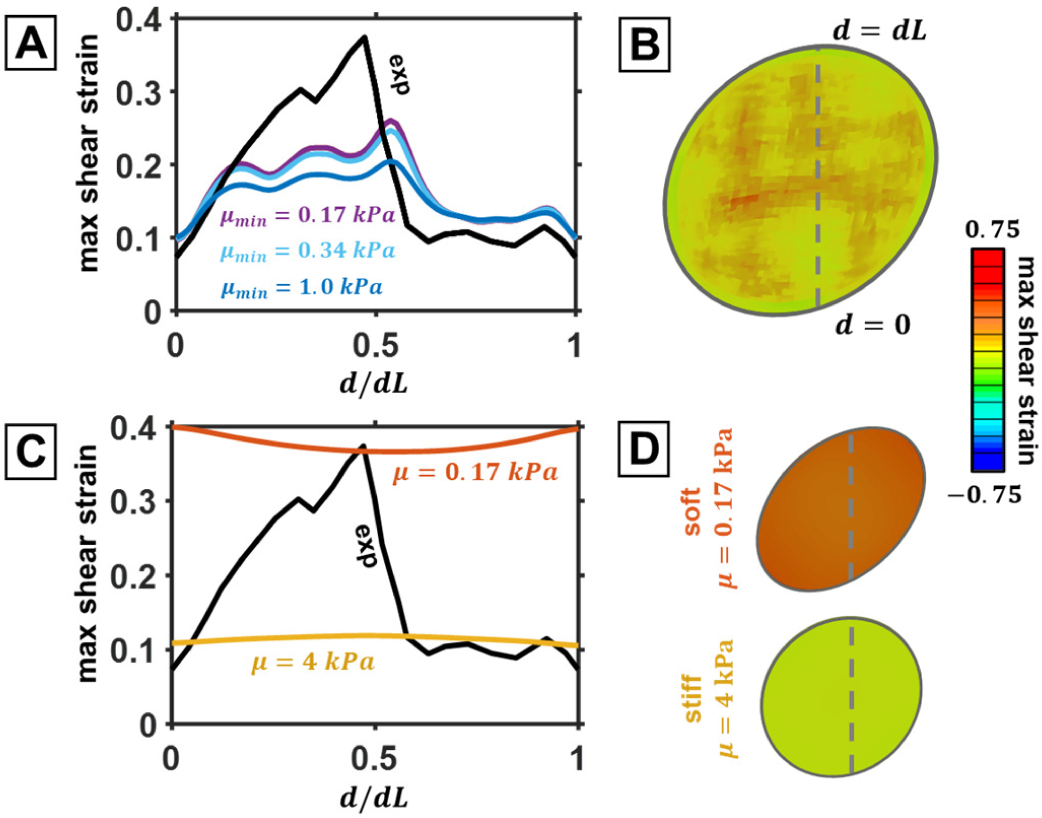
(A) Experimental and predicted shear strain along nucleus section following deformation of a heterogeneously stiff nucleus with a linear distribution of shear moduli, and (B) associated contour plot across nuclear mid-section; (C) Experimental and predicted shear strain along nucleus section following deformation of a homogeneous nucleus (uniform stiffness), and (D) associated contour plot across nuclear mid-section.

## Discussion

This study presents the first computational investigation of the in-situ deformation of a heterogeneous cell nucleus. A novel methodology is developed to accurately reconstruct a three-dimensional finite element spatially heterogeneous model of a cell nucleus from confocal microscopy z-stack images of nuclei stained for nucleus DNA. The relationship between spatially heterogeneous distributions of microscope imaging-derived greyscale values, shear stiffness and resultant shear strain is explored. We incorporate the reconstructed heterogeneous nucleus into a model of a chondrocyte embedded in a PCM and cartilage ECM, based on the micromechanical RVE approach of Dowling et al. (28). Shear loading is applied to the RVE to simulate the experiments of Henderson et al. (26), and computed distributions of shear strain in the heterogeneous nucleus are compared to experimental measurements.

Simulations suggest that the nucleus is highly heterogeneous in terms of its mechanical behaviour with a sigmoidal relationship between experimentally measure greyscale values and corresponding local shear moduli. Three distinct phases are identified within the nucleus: a low stiffness phase (0.17 *kPa* ≤ *μ*_*n*_ ≤ 0.63 *kPa*) corresponding to mRNA rich interchromatin regions; an intermediate stiffness phase (1.48 *kPa* ≤ *μ*_*n*_ ≤ 2.7 *kPa*) corresponding to euchromatin; a high stiffness phase (3.58 *kPa* ≤ *μ*_*n*_ ≤ 4.0 *kPa*) corresponding to heterochromatin. Simulations suggest that disruption of the nucleus envelope as a result of lamin A/C depletion significantly increases nucleus strain in regions of low DNA concentration. The experimental-computational study of Dowling et al. (27) revealed that contractility of the actin cytoskeleton is the key determinant of the response of chondrocytes to externally applied shear deformation. Our simulations suggest that increased cell contractility (as associated with de-differentiation of chondrocytes to a fibroblast-like phenotype) significantly elevates the heterogeneous intra-nuclear strains. As nuclear deformation has been linked to gene expression, alterations in cellular force generation will thus have downstream implications for cell function (35).

Our analysis suggests that the stiffest region of the nucleus (with highest DNA concentration and greyscale value) has a shear modulus that is 23 times higher than that of the most compliant region (with lowest DNA concentration and greyscale value). This pronounced difference in stiffness may be explained in terms of molecular structures of chromosomal DNA and mRNA molecules within the nucleus, given that Henderson et al. (26) observed that regions of low DNA have a high RNA concentration. DNA molecules in eukaryotic cells are thin polymers of length 3-6 cm (36), that must be compacted to fit in the finite volume of a cell nucleus (typically of diameter is 5-10 μm). Thus, the DNA molecule is envisioned as an extremely long thin string of moderate elasticity that is bent into the configurations required for packaging, and, when associated with histones, this compacted complex forms chromatin (37). Moreover, DNA bending is determined by its sequence, which reduces the degrees of bending freedom, and therefore the sequence constrains the number of possible packaging configurations. It is argued that this increases the overall stiffness of the molecule (37). On the other hand, messenger RNA (mRNA) are much shorter molecules (of length ∼300 nm). In the interchromatin of cell nuclei, pre-mRNA molecules are spliced into mature mRNA, and when associated to proteins and non-coding RNA they form complexes known as spliceosomes. Pre-mRNA is not significantly different to mRNA in size, and the average size of spliceosomes is ∼20nm (38). The significantly lower levels of folding in mRNA and spliceosome molecules, relative to DNA, results in a dramatically increased deformability of mRNA regions within the nucleus, as revealed by our heterogeneous nucleus model. Therefore, we suggest that interchromatin regions (rich in mRNA) correspond to the low stiffness regions (LR) identified by our heterogeneous nucleus model, and chromosomic DNA space correspond to the intermediate (IR) and high stiffness (HR) regions. We suggest that the IR and HR regions correspond to euchromatin and heterochromatin, respectively, which are the two main packaging arrangements/ conformations present on each functional chromosome. Euchromatin contains expressed genes and is packaged less densely in order to allow space for the formation of transcription complexes. Gene expression does not occur when chromatin is in the highly densely packed heterochromatin conformation. Moreover, euchromatin is more predominant than heterochromatin in human chromosomes (39), which is consistent with our finding that IR and HR regions comprise of ∼31% and ∼12% of the nucleus volume. Our results also show that the nucleus envelope strongly influences the strain in the nucleus (Fig 4A). This is supported by recent experiments that report chromatin is bound to the lamina of nuclear envelope and therefore influences chromatin deformation (40). Our model predicts that chondrocyte de-differentiation to a fibroblast-like phenotype significantly elevates the heterogeneous intra-nuclear strains. Such a change in deformation of mRNA/ euchromatin/heterochromatin may play a role in altered gene expression and synthesis of type-I collage instead of type-II collagen in late stages of diseased osteoarthritic tissue (8).

The accurate prediction of nucleus deformation is a key step in understanding mechanotransduction. Previous studies have uncovered a link between nucleus deformation and regulation of type II collagen, as well as gene expression (41-45), possibly associated with the reshaping of nuclear lamina and alterations in chromatin distribution (46-48). Using a geometrically accurate 3D model of osteocytes, Verbruggen and others (49) demonstrated that the cell experiences localised strain amplifications due to ECM projections and the presence of a PCM. It was proposed that these strain amplifications are an important mechanism in osteocyte mechanobiology (50). The computational investigation of the current study indicates that nucleus heterogeneity could also be a means by which strain amplification mechanisms promote mechanotransduction.

Our simulations suggest that the range of shear moduli within the nucleus span two orders of magnitude, resulting in peak nucleus strains in localised regions of low DNA concentration that are approximately five times higher than the strains in the ECM. This large range of shear moduli within a single nucleus may explain the many conflicting values in the literature of nucleus stiffness based on the assumption of homogeneous material behaviour. Previous studies on endothelial cells ((10, 16), implementing inverse finite element analysis of experimental unconfined compression tests to calibrate the nucleus mechanical properties assuming material inhomogeneity, report near incompressible behaviour with shear moduli ranging from ∼1 to ∼3 kPa. In contrast, Reynolds et al. (20) developed an experimental methodology to perform micropipette aspiration on spread adhered endothelial cells, including imaging of stress fibres and nucleus. Finite element simulation of experiments using an active stress fibre model resulted in the prediction of a nucleus shear modulus of only 0.07 kPa, again with near incompressibility being observed. Similarly, simulations of micropipette aspiration of isolated chondrocyte nuclei by (21) report a highly compliant homogeneous nucleoplasm with a shear modulus of 0.008 kPa surrounded by stiff lamina and membrane regions in order to replicate micropipette aspiration experimental results. Pipette aspiration studies by Pajerowski et al. (24), Deguchi et al. (15), Guilak et al. (9) and Zhao et al. (19) also report shear moduli lower than 1 kPa for homogeneous nuclei. The significant differences in reported shear moduli, spanning three orders of magnitude, between compression experiments and micropipette aspiration experiments suggests that the assumption of a homogeneous nucleus cannot accurately replicate observed responses to diverse loading using a unique set of material properties. The capability of our highly heterogeneous nucleus to accurately predict experimentally observed nucleus deformations for both compression and micropipette aspiration models should be investigated in future implementations of the model. We speculate that high DNA regions with high shear modulus may contribute strongly to nucleus resistance to parallel plate induced compressive axial deformation, whereas the presence of abundant regions of low shear modulus regions with low DNA concentration would permit significant isochoric deformations during micropipette aspiration.

A potential limitation in the current study is that the nucleus is assumed to be hyperelastic. The deformation applied by Henderson et al. (26) is monotonic and DNA distribution is observed at two time points, once before and once after applied shear deformation. 3D imaging at multiple time-points would provide insight into viscoelasticity and active contractility transience (51). Alternatively, imposing the applied tissue shear at different loading rates could be considered. Several studies suggest that the cell nucleus is elastic with fully recoverable deformation following the application of moderate to large deformations (18, 22, 23). However, Pajerowski et al. (24) report that permanent viscoplastic deformation of the nucleus can occur following knockout of lamin A/C (25) report rupture of the lamin following cell migration through narrow channels. Such inelastic deformation could be investigated in future extensions of this work by imaging nuclei following load removal. Further exploration of the role of the cell cytoskeleton on nucleus deformation should consider the role of cell cytoskeletal free energy in the homeostatic configuration of chondrocytes in-situ (52, 53).

## Acknowledgements

Funding was provided by Science Foundation Ireland grant no. 18/ERCD/5481. The authors also acknowledge the Irish Centre for High-End Computing (ICHEC) for provision of computational facilities and support.

## References

1. Song, Y., J. Soto, B. Chen, L. Yang, and S. Li. 2020. Cell engineering: Biophysical regulation of the nucleus. Biomaterials 234:119743.

2. Bufi, N., M. Saitakis, S. Dogniaux, O. Buschinger, A. Bohineust, A. Richert, M. Maurin, C. Hivroz, and A. Asnacios. 2015. Human Primary Immune Cells Exhibit Distinct Mechanical Properties that Are Modified by Inflammation. Biophysical journal 108(9):2181–2190.

3. Shin, J.-W., K. R. Spinler, J. Swift, J. A. Chasis, N. Mohandas, and D. E. Discher. 2013. Lamins regulate cell trafficking and lineage maturation of adult human hematopoietic cells. Proc Natl Acad Sci U S A 110(47):18892–18897.

4. Irianto, J., C. R. Pfeifer, I. L. Ivanovska, J. Swift, and D. E. Discher. 2016. Nuclear Lamins in Cancer. Cellular and Molecular Bioengineering 9(2):258–267.

5. Cao, X., E. Moeendarbary, P. Isermann, Patricia M. Davidson, X. Wang, Michelle B. Chen, Anya K. Burkart, J. Lammerding, Roger D. Kamm, and Vivek B. Shenoy. 2016. A Chemomechanical Model for Nuclear Morphology and Stresses during Cell Transendothelial Migration. Biophysical journal 111(7):1541–1552.

6. Buxboim, A., J. Irianto, J. Swift, A. Athirasala, J. W. Shin, F. Rehfeldt, and D. E. Discher. 2017. Coordinated increase of nuclear tension and lamin-A with matrix stiffness outcompetes lamin-B receptor that favors soft tissue phenotypes. Molecular biology of the cell 28(23):3333–3348.

7. McBride, S. H., and M. L. Knothe Tate. 2008. Modulation of Stem Cell Shape and Fate A: The Role of Density and Seeding Protocol on Nucleus Shape and Gene Expression. Tissue Engineering Part A 14(9):1561–1572.

8. Charlier, E., C. Deroyer, F. Ciregia, O. Malaise, S. Neuville, Z. Plener, M. Malaise, and D. de Seny. 2019. Chondrocyte dedifferentiation and osteoarthritis (OA). Biochemical pharmacology 165:49–65.

9. Guilak, F., J. R. Tedrow, and R. Burgkart. 2000. Viscoelastic properties of the cell nucleus. Biochem Biophys Res Commun 269(3):781–786.

10. Caille, N., O. Thoumine, Y. Tardy, and J.-J. Meister. 2002. Contribution of the nucleus to the mechanical properties of endothelial cells. J Biomech 35(2):177–187.

11. Caille, N., Y. Tardy, and J. J. Meister. 1998. Assessment of Strain Field in Endothelial Cells Subjected to Uniaxial Deformation of Their Substrate. Annals of Biomedical Engineering 26(3):409–416.

12. Maniotis, A. J., C. S. Chen, and D. E. Ingber. 1997. Demonstration of mechanical connections between integrins, cytoskeletal filaments, and nucleoplasm that stabilize nuclear structure. Proc Natl Acad Sci USA 94(3):849–854.

13. Jean, R. P., C. S. Chen, and A. A. Spector. 2005. Finite-element analysis of the adhesion-cytoskeleton-nucleus mechanotransduction pathway during endothelial cell rounding: axisymmetric model. J Biomech Eng 127(4):594–600.

14. Ferko, M. C., A. Bhatnagar, M. B. Garcia, and P. J. Butler. 2007. Finite-element stress analysis of a multicomponent model of sheared and focally-adhered endothelial cells. Ann Biomed Eng 35(2):208–223.

15. Deguchi, S., M. Yano, K. Hashimoto, H. Fukamachi, S. Washio, and K. Tsujioka. 2007. Assessment of the mechanical properties of the nucleus inside a spherical endothelial cell based on microtensile testing. Journal of Mechanics of Materials and Structures 2(6):1087–1102.

16. Ofek, G., R. M. Natoli, and K. A. Athanasiou. 2009. In situ mechanical properties of the chondrocyte cytoplasm and nucleus. J Biomech 42(7):873–877.

17. Knight, M. M., J. van de Breevaart Bravenboer, D. A. Lee, G. J. V. M. van Osch, H. Weinans, and D. L. Bader. 2002. Cell and nucleus deformation in compressed chondrocyte–alginate constructs: temporal changes and calculation of cell modulus. Biochimica et Biophysica Acta (BBA) - General Subjects 1570(1):1–8.

18. Caille, N., O. Thoumine, Y. Tardy, and J.-J. Meister. 2002. Contribution of the nucleus to the mechanical properties of endothelial cells. Journal of biomechanics 35(2):177–187.

19. Zhao, R., K. Wyss, and C. A. Simmons. 2009. Comparison of analytical and inverse finite element approaches to estimate cell viscoelastic properties by micropipette aspiration. Journal of biomechanics 42(16):2768–2773.

20. Reynolds, N. H., W. Ronan, E. P. Dowling, P. Owens, R. M. McMeeking, and J. P. McGarry. 2014. On the role of the actin cytoskeleton and nucleus in the biomechanical response of spread cells. Biomaterials 35(13):4015–4025.

21. Vaziri, A., and M. R. Mofrad. 2007. Mechanics and deformation of the nucleus in micropipette aspiration experiment. J Biomech 40(9):2053–2062.

22. Hu, S., R. Wang, C. M. Tsang, S. W. Tsao, D. Sun, and R. H. W. Lam. 2018. Revealing elasticity of largely deformed cells flowing along confining microchannels. RSC Advances 8(2):1030-1038. 10.1039/C7RA10750A.

23. Weafer, P. P., W. Ronan, S. P. Jarvis, and J. P. McGarry. 2013. Experimental and Computational Investigation of the Role of Stress Fiber Contractility in the Resistance of Osteoblasts to Compression. Bulletin of Mathematical Biology 75(8):1284–1303.

24. Pajerowski, J. D., K. N. Dahl, F. L. Zhong, P. J. Sammak, and D. E. Discher. 2007. Physical plasticity of the nucleus in stem cell differentiation. Proc Natl Acad Sci U S A 104(40):15619–15624.

25. Thiam, H.-R., P. Vargas, N. Carpi, C. L. Crespo, M. Raab, E. Terriac, M. C. King, J. Jacobelli, A. S. Alberts, T. Stradal, A.-M. Lennon-Dumenil, and M. Piel. 2016. Perinuclear Arp2/3-driven actin polymerization enables nuclear deformation to facilitate cell migration through complex environments. Nature Communications 7(1):10997.

26. Henderson, J. T., G. Shannon, A. I. Veress, and C. P. Neu. 2013. Direct measurement of intranuclear strain distributions and RNA synthesis in single cells embedded within native tissue. Biophysical journal 105(10):2252–2261.

27. Dowling, E. P., W. Ronan, G. Ofek, V. S. Deshpande, R. M. McMeeking, K. A. Athanasiou, and J. P. McGarry. 2012. The effect of remodelling and contractility of the actin cytoskeleton on the shear resistance of single cells: a computational and experimental investigation. Journal of The Royal Society Interface 9(77):3469–3479.

28. Dowling, E. P., W. Ronan, and J. P. McGarry. 2013. Computational investigation of in situ chondrocyte deformation and actin cytoskeleton remodelling under physiological loading. Acta Biomaterialia 9(4):5943–5955.

29. Nolan, D. R., A. L. Gower, M. Destrade, R. W. Ogden, and J. P. McGarry. 2014. A robust anisotropic hyperelastic formulation for the modelling of soft tissue. Journal of the mechanical behavior of biomedical materials 39:48–60.

30. Deshpande, V. S., R. M. McMeeking, and A. G. Evans. 2006. A bio-chemo-mechanical model for cell contractility. Proceedings of the National Academy of Sciences 103(38):14015–14020.

31. McGarry, J. P. 2009. Characterization of cell mechanical properties by computational modeling of parallel plate compression. Ann Biomed Eng 37(11):2317–2325.

32. Ronan, W., V. S. Deshpande, R. M. McMeeking, and J. P. McGarry. 2014. Cellular contractility and substrate elasticity: a numerical investigation of the actin cytoskeleton and cell adhesion. Biomechanics and Modeling in Mechanobiology 13(2):417–435.

33. McGarry, J. P., and P. E. McHugh. 2008. Modelling of in vitro chondrocyte detachment. Journal of the Mechanics and Physics of Solids 56(4):1554–1565.

34. Dowling, E. P., and J. P. McGarry. 2014. Influence of Spreading and Contractility on Cell Detachment. Annals of Biomedical Engineering 42(5):1037–1048.

35. Alisafaei, F., D. S. Jokhun, G. V. Shivashankar, and V. B. Shenoy. 2019. Regulation of nuclear architecture, mechanics, and nucleocytoplasmic shuttling of epigenetic factors by cell geometric constraints. Proceedings of the National Academy of Sciences 116(27):13200–13209.

36. Piovesan, A., M. C. Pelleri, F. Antonaros, P. Strippoli, M. Caracausi, and L. Vitale. 2019. On the length, weight and GC content of the human genome. BMC Res Notes 12(1):106–106.

37. Vivante, A., I. Bronshtein, and Y. Garini. 2020. Chromatin Viscoelasticity Measured by Local Dynamic Analysis. Biophysical journal 118(9):2258–2267.

38. Sperling, J., M. Azubel, and R. Sperling. 2008. Structure and function of the Pre-mRNA splicing machine. Structure (London, England : 1993) 16(11):1605–1615.

39. International Human Genome Sequencing, C. 2004. Finishing the euchromatic sequence of the human genome. Nature 431(7011):931–945.

40. Stephens, A. D., E. J. Banigan, and J. F. Marko. 2019. Chromatin’s physical properties shape the nucleus and its functions. Current opinion in cell biology 58:76–84.

41. Roca-Cusachs, P., J. Alcaraz, R. Sunyer, J. Samitier, R. Farre, and D. Navajas. 2008. Micropatterning of single endothelial cell shape reveals a tight coupling between nuclear volume in G1 and proliferation. Biophys J 94(12):4984–4995.

42. Thomas, C. H., J. H. Collier, C. S. Sfeir, and K. E. Healy. 2002. Engineering gene expression and protein synthesis by modulation of nuclear shape. Proc Natl Acad Sci USA 99(4):1972–1977.

43. Campbell, J. J., E. J. Blain, T. T. Chowdhury, and M. M. Knight. 2007. Loading alters actin dynamics and up-regulates cofilin gene expression in chondrocytes. Biochem Biophys Res Commun 361(2):329–334.

44. Buschmann, M. D., E. B. Hunziker, Y. J. Kim, and A. J. Grodzinsky. 1996. Altered aggrecan synthesis correlates with cell and nucleus structure in statically compressed cartilage. J Cell Sci 109(Pt 2):499–508.

45. Shieh, A. C., and K. A. Athanasiou. 2007. Dynamic compression of single cells. Osteoarthritis Cartilage 15(3):328–334.

46. Dahl, K. N., A. J. Ribeiro, and J. Lammerding. 2008. Nuclear shape, mechanics, and mechanotransduction. Circ Res 102(11):1307–1318.

47. Rowat, A. C., J. Lammerding, H. Herrmann, and U. Aebi. 2008. Towards an integrated understanding of the structure and mechanics of the cell nucleus. Bioessays 30(3):226–236.

48. Tsukamoto, T., N. Hashiguchi, S. M. Janicki, T. Tumbar, A. S. Belmont, and D. L. Spector. 2000. Visualization of gene activity in living cells. Nat Cell Biol 2(12):871–878.

49. Verbruggen, S. W., T. J. Vaughan, and L. M. McNamara. 2012. Strain amplification in bone mechanobiology: a computational investigation of the in vivo mechanics of osteocytes. J R Soc Interface 9(75):2735–2744. Research Support, Non-U.S. Gov’t.

50. Adachi, T., K. Sato, N. Higashi, Y. Tomita, and M. Hojo. 2008. Simultaneous observation of calcium signaling response and membrane deformation due to localized mechanical stimulus in single osteoblast-like cells. Journal of the Mechanical Behavior of Biomedical Materials 1(1):43–50.

51. McEvoy, E., V. S. Deshpande, and P. McGarry. 2019. Transient active force generation and stress fibre remodelling in cells under cyclic loading. Biomechanics and Modeling in Mechanobiology 18(4):921–937.

52. McEvoy, E., V. S. Deshpande, and P. McGarry. 2017. Free energy analysis of cell spreading. Journal of the mechanical behavior of biomedical materials 74:283–295.

53. McEvoy, E., S. S. Shishvan, V. S. Deshpande, and J. P. McGarry. 2018. Thermodynamic Modeling of the Statistics of Cell Spreading on Ligand-Coated Elastic Substrates. Biophysical journal 115(12):2451–2460.

